# A Versatile Microfluidic Device for High-throughput Combinatorial Drug Screening

**DOI:** 10.64898/2026.09.09.750366

**Authors:** Saumya Jaiswal, Mallikarjun PVN Reddy, Debjani Paul, Prasanna Gandhi, Abhijit Majumder

**Affiliations:** Indian Institute of Technology Bombay, Department of Chemical Engineering, Mumbai, India; Indian Institute of Technology Bombay, Department of Mechanical Engineering, Mumbai, India; Indian Institute of Technology Bombay, Department of Biosciences and Bioengineering Engineering, Mumbai, India; Johns Hopkins University, Department of Chemical and Biomolecular Engineering, Baltimore, United States

**Keywords:** Drug screening, microfluidic device, concentration gradient generator (CGG), combinatorial drug screening

## Abstract

Combination therapies can improve anticancer efficacy, but identifying effective drug pairs and dose combinations requires systematic exploration of multidimensional concentration spaces. Here, we extend a diffusion-driven, flowless microfluidic concentration gradient generator (CGG) to enable quantitative combinatorial drug screening without external pumps or continuous flow. The platform comprises a 5 × 5 array of interconnected culture nodes coupled to four peripheral reservoirs, in which overlapping diffusion fields generate spatially defined single- and multidrug exposures. Computational modelling was used to assign local drug concentrations to individual nodes, enabling direct correlation of the predicted exposure landscape with cellular response. Using MCF-7 breast cancer cells and 5-fluorouracil (5-FU) and doxorubicin (DOX) as model therapeutics, the platform resolved dependent single-agent responses, yielding IC_50_ values of 3.46 μM for 5-FU and 1.22 μM for DOX. Combinatorial loading generated 25 spatially defined 5-FU DOX concentration pairs within a single device, which were resolved into two-dimensional concentration-response landscapes. Bliss independence analysis revealed concentration-specific drug interactions, with synergy predominating at low-to-intermediate concentrations and a transition towards additive and antagonistic responses at higher exposures. These findings establish a pump-free microfluidic strategy that integrates computational concentration mapping with spatially resolved pharmacological analysis to identify effective drug-combination windows within a single platform.

## 1. Introduction

Combination therapy is widely used in cancer treatment to enhance therapeutic efficacy and overcome resistance by simultaneously perturbing complementary cellular pathways [1], [2]. Appropriately selected drug combinations can also reduce the doses required for individual agents, potentially improving therapeutic selectivity and limiting dose-associated toxicity [3]. However, drug interactions are inherently concentration dependent: the same pair of compounds can produce different responses depending on their absolute concentrations and relative exposure. Identifying effective combinations therefore requires interrogation of a multidimensional dose space rather than evaluation of a limited number of arbitrarily selected drug pairs [4].

Conventional combination screening typically relies on multiwell assays in which individual drug concentrations and ratios are prepared by serial dilution and distributed across separate wells. Although robust and widely adopted, this approach rapidly increases the number of experimental conditions, cells and reagents required as the concentration space expands [5], [6]. Microfluidic platforms provide an attractive alternative by generating multiple chemical environments within miniaturized culture systems, enabling concentration gradients and drug mixtures to be evaluated with substantially reduced sample requirements [7], [8], [9]. Recent advances have extended these platforms to combinatorial drug screening by generating multidrug concentration landscapes and mixture combinations for high-throughput evaluation of therapeutic responses. These systems have enabled screening of multiple drug concentrations, ratios, and patient-derived models, highlighting the potential of microfluidics for precision medicine [10]. However, most existing platforms rely on syringe pumps, pressure-driven flow, pneumatic control, multilayer architectures, or complex fluidic networks, which can increase fabrication complexity and restrict routine implementation [11]. Consequently, there remains a need for simple, pump-free microfluidic platforms capable of generating stable multidrug concentration gradients under static culture conditions while remaining compatible with long-term cell culture and downstream biological analyses.

We previously developed a flowless, diffusion-driven microfluidic platform capable of generating stable concentration gradients for single-drug screening without external pumping or continuous reservoir replenishment [12]. Subsequent theoretical and computational studies established the design principles governing long-term gradient stability and extended this framework to the generation of predictable multi-solute concentration landscapes [13]. Despite this engineering framework, the multi-solute configuration had not been experimentally translated into combinatorial drug screening, leaving unresolved whether spatially generated drug combinations could be quantitatively linked to cellular response and drug interaction.

Here, we extend this static diffusion-based microfluidic platform to enable quantitative combinatorial drug screening by generating orthogonal concentration gradients of multiple therapeutics within a single device. Using MCF-7 breast cancer cells and 5-fluorouracil (5-FU) and doxorubicin (DOX) as model therapeutics, we couple computationally predicted local drug concentrations with spatially resolved cell-survival measurements to derive single-agent dose-response relationships and multidimensional combination-response landscapes. Bliss independence analysis further resolves concentration-specific synergistic, additive and antagonistic interactions across the generated drug space. This work advances the flowless CGG from a computationally predicted multi-solute framework to an experimentally validated platform for quantitative combinatorial drug screening.

## 2. Results

### 2.1 Device design and fabrication

The microfluidic platform was fabricated by CNC micromilling-assisted soft lithography to generate a PDMS device comprising four reservoirs interconnected by an array of hemispherical culture nodes and rectangular diffusion channels. The fabricated device faithfully reproduced the designed microarchitecture and formed a leak-free PDMS-glass assembly following plasma bonding, enabling prolonged diffusion-based experiments and cell culture. Building upon our previously validated gradient-generation platform [13], the present device was adapted for combinatorial drug screening by integrating orthogonal drug reservoirs that generate multiple concentration combinations across the node array. The resulting architecture provides an optically transparent and reproducible platform compatible with fluorescence imaging, live-cell assays, and downstream immunostaining.

### 2.2 Uniform cell seeding across the microfluidic node array

Uniform cell seeding across the culture nodes is essential for reliable quantification of drug-induced cytotoxicity, as variations in the initial cell number can confound the interpretation of concentration-dependent responses. To optimize cell loading, MCF-7 cells were seeded at different cell densities, and the resulting distribution was quantified after cell attachment.

The hemispherical node architecture facilitated efficient cell capture, resulting in homogeneous cell distribution throughout the microfluidic network (Fig. 2A). Increasing the seeding density progressively increased the number of cells retained within each node, while reducing the variability in cell occupancy. Among the tested conditions, a seeding density of 0.75 × 10^5^ cells per device produced the most uniform distribution, with an average of 150 ± 30 cells per node, corresponding to approximately 3,750 cells retained across the 25-node array (Fig. 2B). This seeding condition provided sufficient cell numbers for downstream viability analysis while maintaining minimal node-to-node variation.

**Figure 1.**
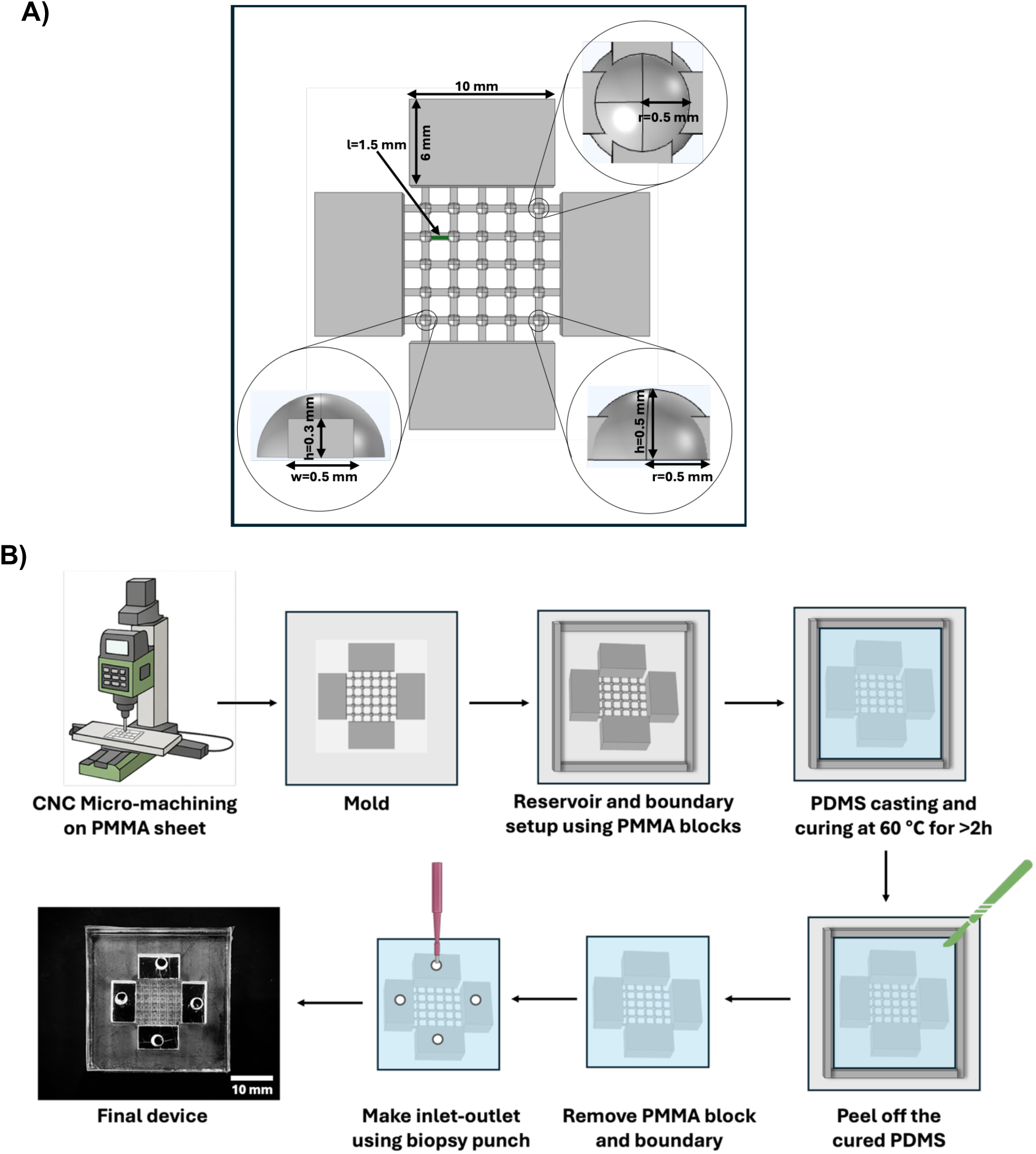
**A)** Design and **B)** fabrication protocol of the combinatorial drug screening microfluidic device.

**Figure 2.**
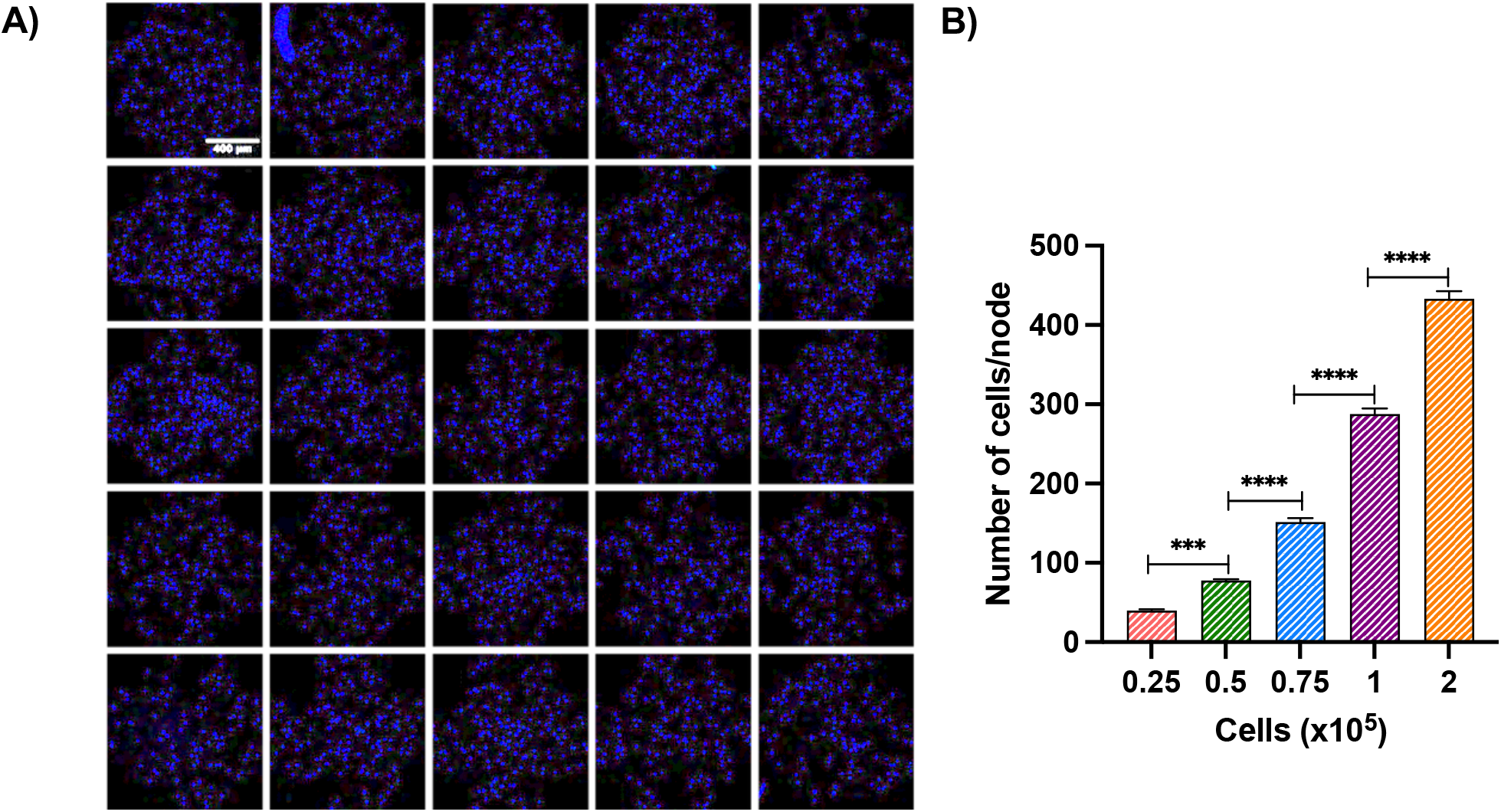
Optimization of uniform cell seeding in the microfluidic device. **(A)** Representative Hoechst stained image showing the uniform distribution of MCF-7 cells across the 25 nodes following seeding at an optimized density of 0.75 × 10^5^ cells per device. **(B)** Quantification of the average number of cells per node at different seeding densities. Data are presented as mean ± SD. Scale bar- 400μm

### 2.3 Combinatorial drug testing in microfluidic device

We next examined how the spatially resolved 5-FU and DOX concentration profiles translated into single-agent and combinatorial responses in MCF-7 cells. The 5 × 5 CGG comprised 25 interconnected cell-culture nodes coupled to four peripheral reservoirs, enabling diffusion-mediated exposure of cells to spatially varying drug concentrations (Fig. 3A). COMSOL simulations were first used to determine the local concentrations established under each drug-loading condition (Fig. 3B-E). Single-agent loading of 5-FU (5 µM) (Fig. 3B) or DOX (1.5 µM) (Fig. 3C) produced graded concentration profiles across the network. In contrast, introducing the two drugs from different reservoirs generated intersecting diffusion fields, assigning a distinct 5-FU-DOX concentration pair to each culture node (Fig. 3D, E). Changing the reservoir-loading scheme altered both the magnitude and distribution of these concentration pairs, thereby extending the range of combinations sampled within the same device geometry.

**Figure 3.**
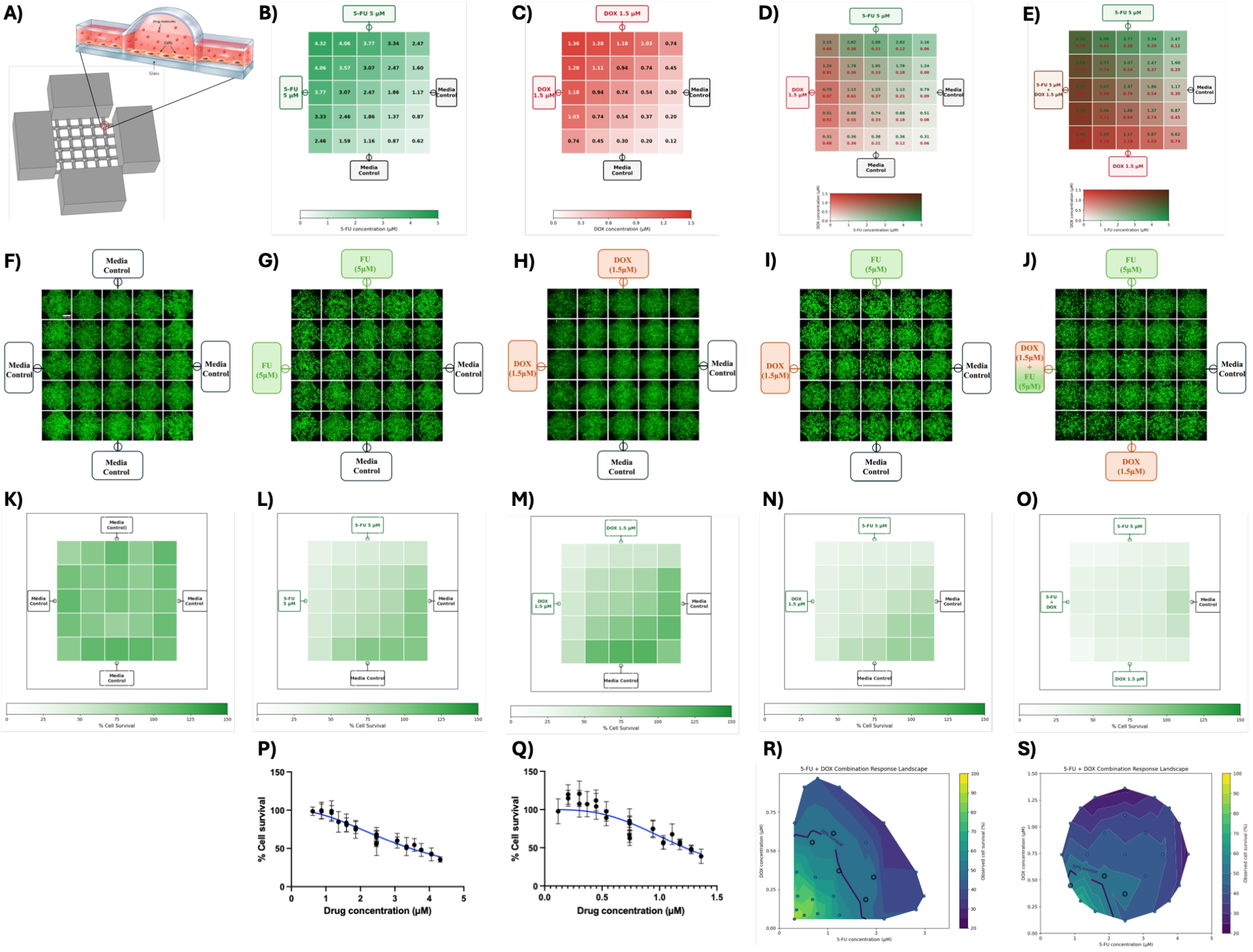
Experimental validation of a flowless concentration gradient generator for single and combinatorial drug screening. **(A)** Schematic of the four-reservoir, 5 × 5 flowless CGG, showing cell localization within the interconnected culture nodes and channels. **(B-E)** COMSOL-predicted spatial concentration profiles for the different drug-loading configurations: **(B)** 5-FU single-drug gradient, **(C)** DOX single-drug gradient, **(D)** two-reservoir 5-FU + DOX combination, and **(E)** three-reservoir 5-FU + DOX combination. **(F-J)** Representative fluorescence images of MCF-7 cells within the CGG following treatment under the corresponding control, single-drug, and combinatorial drug conditions. Scale bar-400μm **(K-O)** Spatial maps of percentage cell survival across the 25 culture nodes for the corresponding treatment conditions. **(P-Q)** Concentration-response curves for single-agent **5-FU** and **DOX**, respectively. **(R-S)** Two-dimensional concentration-response landscapes showing experimentally observed MCF-7 cell survival across the 5-FU–DOX concentration space for the two-reservoir and three-reservoir combinatorial configurations, respectively.

These spatial differences in drug exposure were reflected in the experimental response after 48 h treatment (Fig. 3F-O). Untreated MCF-7 cells maintained high survival across the network, whereas exposure to either drug alone resulted in progressively lower survival with increasing local concentration. Relating the experimentally measured survival at each node to its corresponding simulated concentration generated well-defined single-agent dose-response relationships (Fig. 3P, Q). The fitted curves yielded an IC_50_ of 3.46 µM for 5-FU and 1.22 µM for DOX, with Hill coefficients of 2.05 and 3.12, respectively.

Combination treatment produced a substantially broader pattern of cellular responses than single-agent gradient (Fig. 3N, O). In the first configuration, the 25 nodes encompassed 0.31-2.98 µM 5-FU and 0.059-0.97 µM DOX, whereas the second configuration extended the exposure space to 0.62-4.31 µM 5-FU and 0.12-1.36 µM DOX. Each of the 25 culture nodes therefore represented a defined combination of the two drugs rather than a replicate condition.

The resulting spatial survival maps showed pronounced variation across the network, which was resolved as two-dimensional concentration–response landscapes by plotting survival against the corresponding local concentrations of both agents (Fig. 3R, S). The first configuration (Fig 3R) predominantly sampled concentrations below the respective single-agent IC_50_ values, yet substantial reductions in survival were observed at several combination conditions. Extending the concentration range in the second configuration (Fig. 3S) produced a broader response surface, with progressively lower survival as the concentrations of both agents increased. Thus, the cellular response was determined by the specific 5-FU-DOX concentration pair rather than by spatial position alone.

Collectively, these results show that the same flowless CGG can resolve conventional single-agent pharmacological parameters and multidimensional combination responses from spatial concentration gradients. The pronounced responses observed for several combinations at concentrations below the respective single-agent IC_50_ values suggested that the effect of combined exposure could not be interpreted from single-agent potency alone. The heterogeneous responses observed across this concentration space provided the basis for subsequently determining whether individual drug combinations produced synergistic, additive or antagonistic interactions.

### 2.4 Synergy scoring

To define the interaction between 5-FU and DOX across the concentration space generated within the CGG, the observed combination responses were evaluated using the Bliss independence model (Fig. 4). In the first drug-loading configuration (Fig. 4A), all tested concentration pairs showed synergistic interactions, with the strongest synergy occurring within the lower concentration range. The interaction weakened progressively with increasing drug exposure, as the Bliss-derived index shifted towards unity. Expanding the concentration range in the second configuration (Fig. 4B) revealed a clear transition in drug interaction, synergy was retained across low-to-intermediate concentration combinations, whereas higher concentrations increasingly produced additive and, at selected combinations, antagonistic responses. These findings show that maximal drug exposure did not correspond to maximal synergy; rather, the cooperative effect of 5-FU and DOX was restricted to defined concentration windows. The spatially resolved interaction profiles therefore demonstrate the ability of the CGG to identify optimal synergistic dose combinations while simultaneously resolving the transition from synergy to additivity and antagonism within a single platform.

**Figure4.**
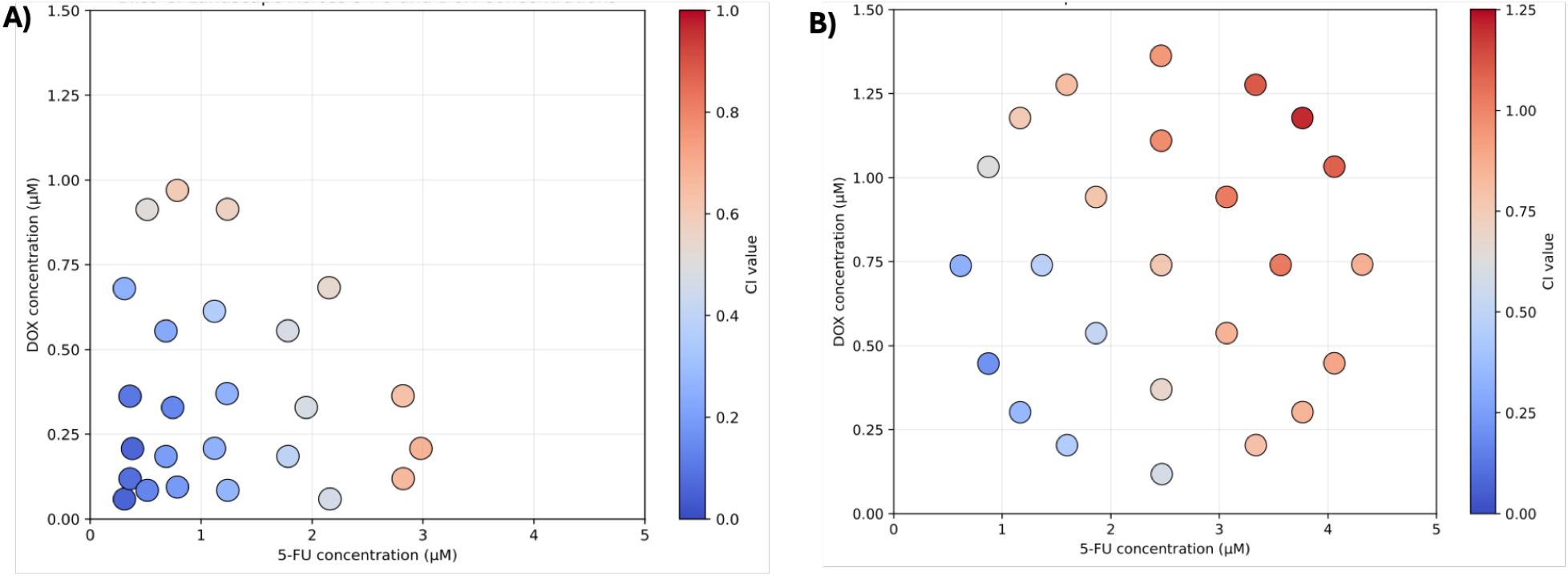
Spatially resolved Bliss analysis reveals distinct synergistic windows for 5-FU and DOX combinations. Two-dimensional interaction landscapes obtained for (A) two-reservoir 5-FU + DOX combination, and **(B)** three-reservoir 5-FU + DOX combination. Bliss-derived interaction index values <1, ≈1 and >1 represent synergistic, additive and antagonistic interactions, respectively.

## 3. Materials and Methods

### 3.1 Fabrication of microfluidic device

#### 3.1.1 Template fabrication

In the CNC micro-milling process, the master template’s micro-channel and node network were created. The template was designed using AutoCAD 2023 software to generate computer-aided design (CAD) files. The CAD design was converted to Standard Tessellation Language (STL) format, and SRP player software was then used as a computer-aided manufacturing (CAM) program to convert the STL file into numerical control (NC) programming language (G-code files). These files were run on Roland SRM 20 micro-milling machine at Srinivasan and Nagar Microfactory, IIT Bombay. Polymethyl methacrylate (PMMA) with a 4 mm thickness was served as the substrate for mold fabrication in the micro-milling process. A single-flute square end mill with a diameter of 0.8 mm and a flute length of 2.4 mm were used as the cutting tool. The spindle speed was set at 7,000 rpm, the feed rate at 450 mm/min, and the depth of the cut at 0.1 mm. Before micro-milling, the cutting tool’s tip was zeroed on the top surface of the PMMA sheet using a touchpoint sensor. The finishing step employed a ball mill with a flute diameter of 1 mm and a flute length of 2 mm, with a feed rate of 600 mm/min. No further surface smoothening methods were necessary, making this master mold ready for device fabrication.

#### 3.1.2 Preparation of the microfluidic device

The pattern on the master template is transferred onto polydimethyl siloxane (PDMS, Sylgard 184) using soft lithography. PDMS is a thermally curable elastomer widely used in microfluidics due to its biocompatibility, nontoxicity, and transparency [14]. First, PDMS and curing agent were mixed in a 10:1 ratio and stirred for 5 min. A hollow PMMA rectangle of height 6 mm was stuck on the template using PDMS stamping. This was done to confine the PDMS such that it is centered at the master pattern. PMMA blocks of 15 × 7×3 mm were also stamp-bonded with PDMS at the reservoir locations, and cured for 15-20 min at 60 °C. These serve to mold the reservoir in the template. The mixture was poured on the master mold and then degassed in a vacuum chamber for 30 min and is subsequently cured in a hot air oven at 60 °C for 2 h. The PDMS layers of 6 mm thickness were carefully peeled off from the mold, and the mold can be reused to make more templates.

In the next step, the inlet and the outlet holes were punched centered on both the reservoirs using a 2 mm Biopsy punch. Then, the PDMS surface was cleaned with scotch tape and then bonded onto a glass slide with plasma treatment (Harrick Plasma) for 180 s. Finally, the chip was filled with Collagen type 1 (25μg/ml) and kept overnight at 4 ºC for coating.

### 3.2 Cell culture

The MCF-7 cells (breast cancer cell line) were used for drug testing. The cells were cultured in high-glucose Dulbecco’s modified Eagle medium (DMEM) (HiMedia-AL007A), supplemented with 1% antibiotic-antifungal solution (HiMedia-A002), 1% L-glutamine (Gibco), and 10% fetal bovine serum (FBS) (HiMedia-RM1112) in a T25 flask until it reaches 80% confluence. For trypsinization, cells were rinsed with DPBS (Dulbecco’s phosphate-buffered saline) (HiMedia-TS1006), followed by the addition of 1 ml of trypsin-EDTA 0.05% (Himedia-TCL0033) to a T-25 flask containing cells at 70-80% confluency. The trypsin and cells were then incubated at 37°C for 5 minutes. Then, the trypsin was neutralized with 1 ml of complete media and transferred into a 15 ml falcon. The resulting cell suspension was centrifuged at 1200 rpm for 5 min to obtain a cell pellet. The cell pellet was resuspended in fresh media, and cells were counted using a hemocytometer (Invitrogen).

### 3.3 Cell seeding in the device

The fabricated device was rinsed twice with DPBS and once with media to remove the excess/unbound collagen in the channel. Then, cell suspension with the desired cell density was seeded in the inlet reservoir. Immediately after seeding, the device was tilted approximately 10-12 times from all sides for uniform distribution of cells at the nodes. After ensuring uniform cell distribution among all nodes under a microscope, excess cell suspension in the reservoirs was removed using 2 mm gauge syringe needle with volume capacity of 2 ml. Subsequently, the device was placed in a CO_2_ incubator at 37 °C for 2 h to facilitate cell attachment to the bottom surface of channel. Once the cells attached to the channel surface, the device was filled with media containing 10% FBS, ensuring the absence of air entrapment within the channels.

### 3.4 Drug gradient generation in the device

For drug testing, 0.75 × 10^5^ cells were seeded in five microfluidic devices and cultured in complete media without drugs for 24 h to allow cell attachment and stabilization. After this incubation period, the devices were divided into two groups based on media conditions: test devices (n = 4) and negative control (n = 1). In the test devices, four drug configurations were established: (i) 5-FU in two adjacent reservoirs, (ii) DOX in two adjacent reservoirs, (iii) 5-FU and DOX in adjacent reservoirs, and (iv) a sequential arrangement consisting of 5-FU in one reservoir, a combination of 5-FU and DOX in the adjacent reservoir, and DOX in the next reservoir, with the remaining reservoir serving as a sink reservoir. The negative control device contained only culture media without drug treatment. Cells were exposed to the resulting drug concentration gradients for 48 h, after which cell viability was assessed using a Live-Dead assay.

### 3.5 Live-dead assay in the device

After 48 hours of 5-FU and DOX drug treatment, the media from the test and control devices were carefully removed. Then, fresh media containing Hoechst (4′,6-diamidino-2-phenylindole) (1:3000) (Life Technologies, Cat. No. H3570), Calcein AM (1:2000) (Invitrogen, Cat. No. C3099) and propidium iodide (PI) (Sigma Aldrich, Cat. No. 81845) (1:1000) was added in the devices and incubated at 37°C for 15 min. Images were captured for all the nodes of the test and control devices in the DAPI, GFP and RFP channels of fluorescence microscope (NIKON Ti2 eclipse). Later, we counted the number of nuclei at each node. The following are the calculations performed after counting the cell number:

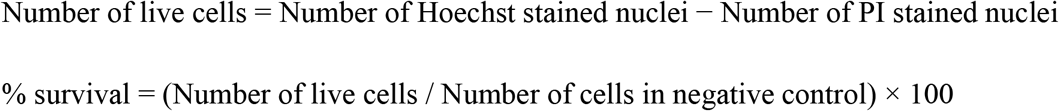

### 3.6 Bliss independence calculations for drug combination effects

Drug interactions between 5-fluorouracil (5-FU) and doxorubicin (DOX) were evaluated using the Bliss independence model. The expected combination effect (*E*_Bliss_) at each concentration pair was calculated from the corresponding single-agent effects as:

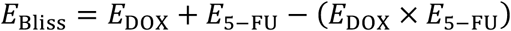

where *E*_Dox_ and *E*_5−FU_ represent the fractional effects of the individual drugs at the corresponding concentrations. Single-agent effects were estimated from the fitted concentration-response curves using the local drug concentrations predicted for each node of the CGG. The predicted Bliss effect was subsequently compared with the experimentally observed combination effect (*E*_Dox+5−FU_) using a combination index

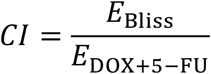

CI values <1, =1 and >1 were classified as synergistic, additive and antagonistic interactions, respectively. The calculated CI values were mapped against the corresponding 5-FU and DOX concentrations to generate two-dimensional interaction landscapes [15].

## 4. Discussion

This study extends a previously established flowless concentration gradient generator from single-drug screening and computational multi-solute prediction to experimental combinatorial drug screening. By integrating COMSOL-derived local concentrations with spatially resolved cellular responses, the platform enabled quantitative assessment of both single-agent and combination treatments within the same microfluidic architecture. The single-agent gradients yielded IC_50_ values of 3.46 μM for 5-FU and 1.22 μM for DOX, demonstrating that spatial concentration gradients can be translated into conventional pharmacological dose-response parameters.

An important feature of the platform is its ability to convert overlapping diffusion gradients into a multidimensional drug-response space. Each of the 25 culture nodes represented a distinct 5-FU-DOX concentration pair, and altering the reservoir-loading configuration shifted the range of combinations sampled without requiring modification of the device geometry. This provides a simple means of exploring multiple drug concentrations and ratios while avoiding the extensive serial dilution and parallel culture conditions required in conventional well-plate screening.

Bliss analysis showed that the interaction between 5-FU and DOX was strongly dependent on concentration. Synergy predominated at low-to-intermediate concentrations, whereas increasing drug exposure shifted the response towards additivity and, at selected combinations, antagonism. Thus, greater cytotoxicity at higher concentrations did not necessarily correspond to greater synergy, emphasizing the need for identifying effective combination windows rather than simply maximizing drug exposure. The ability to resolve these transitions within a single concentration landscape represents a major advantage of spatial gradient-based combination screening.

The current work provides a proof-of-concept using MCF-7 cells and one drug pair, and further validation across additional drugs and cancer models will be required to establish broader applicability. Direct experimental measurement of local therapeutic concentrations would also strengthen the computationally derived exposure maps. However, the pump-free architecture, low operational complexity and integration of concentration prediction with biological response provide a flexible framework for combinatorial screening. Extending the platform to three-dimensional tumor models and patient-derived cells could enable drug-response testing under more physiologically relevant conditions.

## 5. Conclusion

We experimentally established a flowless microfluidic concentration gradient generator for quantitative combinatorial drug screening. By coupling diffusion-driven drug gradients with computational concentration mapping and spatially resolved cellular responses, the platform enabled single-agent dose-response analysis and simultaneous evaluation of multiple 5-FU-DOX concentration combinations within a single device. Bliss analysis further revealed that drug interactions were concentration dependent, with distinct transitions between synergistic, additive and antagonistic responses across the explored concentration space. Overall, these results demonstrate that predictable diffusion fields can be translated into quantitative pharmacological response landscapes without external pumping or complex fluidic control. With further validation across diverse therapeutics and physiologically relevant tumor models, this approach could provide a simple and adaptable framework for identifying favorable combination-dose windows and prioritizing conditions for subsequent preclinical evaluation.

